# ecoTolerance: An R package for Assessing Road and Human Footprint Tolerance in Wildlife Species

**DOI:** 10.64898/2026.02.26.708267

**Authors:** Diego Fernandes Miranda, Lucas Rodriguez Forti

## Abstract

Most wildlife species currently inhabit areas transformed by human activity, a hallmark of the Anthropocene. Habitat alterations caused by the creation of roads and other human-made infrastructures shape the spatial distribution of wildlife species and their interaction with the environment. While some sensitive species disappear, more tolerant ones thrive near humans. Therefore, a streamlined tool to quantify the tolerance of different species to human pressures is useful to conservation, in particular to identify more vulnerable species. Here, we present ecoTolerance, an open-source R package that calculates two complementary, continuous metrics: the Road Tolerance Index (RTI), derived from the distance of each occurrence record to the nearest road, and the Human-Footprint Tolerance Index (HFTI), based on the global human-footprint raster. This package is based on a workflow that includes separate functions and arguments to automate data cleaning, spatial thinning, distance extraction, species-level summarization and map generation. As an applied example of its use and application, we processed 3782 records of five species: *Copaifera langsdorffii* (1407 observations), *Bradypus variegatus* (724), *Sylvilagus brasiliensis* (274), *Boana faber* (1226), and *Boana boans* (151), revealing RTI values that ranged from 0.183 to 0.654 and HFTI values from 0.111 to 0.392. the values of the two indices varied according to the incidence of road kill, as well as the habitat preference of the particular species. These examples demonstrate that ecoTolerance facilitates a rapid and streamlined assessment of species tolerance and vulnerability, providing valuable insights with potential to inform conservation actions.

## 1 INTRODUCTION

Global human population has recently exceeded eight billion (Adam, 2022), and the impact of human activity is evident across much of the planet’s surface. Land-use change driven by anthropogenic processes, such as urbanization and the creation of linear energy and transportation infrastructures, is a key factor in biodiversity loss (Di Marco et al., 2018; Laurance et al., 2009; Piano et al., 2020). As a consequence of human impact, the number of terrestrial vertebrates has declined by 40% between 1970 and 2016 according to the IPBES Global Assessment (Díaz et al., 2019). Human activities lead to deforestation and habitat fragmentation, altering natural environments, reducing connectivity, and negatively affecting wildlife, especially vulnerable species (Fahrig, 2003; Foley et al., 2005). Therefore, human activities affect species distribution (Frans & Liu, 2024). Nevertheless, while many species disappear with increased human presence, others persist or even thrive in highly modified environments (Niederman et al., 2025).

Quantifying the tolerance to or avoidance of anthropogenic environments of particular species is essential for guiding evidence-based conservation strategies and planning land-use and land-cover changes, especially in species-rich regions that are under pressure for development. Some indices have been described to quantify species’ tolerance to human-altered environments from an ecological perspective (Fanelli et al., 2022; Marjakangas et al., 2024). However, many of these indices are not easily transferred among different taxa or are calculated through a common tool among different ecologists. Here we present two indices that can indicate tolerance, use, or avoidance of areas impacted by roads or with high human footprint. One such index is the Road Tolerance Index (RTI), which can be inferred from assessing the spatial distribution of the species relative to a pre-established road network (Dai et al., 2025; González-Bernardo et al., 2023). On the other hand, Human Footprint Tolerance Index (HFTI) evaluates the extent to which a species is found across areas with varying degrees of human modification, including different levels of artificial light at night (ALAN), transportation corridors, built environments, agricultural land and pasture (Mu et al., 2022).

Calculating these indices involves data preparation to reduce bias, buffering, spatial joins, and statistical summarization, which can be time-consuming and prone to significant variation and inconsistencies across studies, especially with the variety of software tools and scripts. Therefore, there is a demand for a streamlined pipeline-processing tool, particularly for large-scale datasets.

R is a programming language (R Core Team, 2025) that stands out as an important platform for ecological research due to its flexibility in statistical computing and extensive ecosystem of packages (Liu et al., 2021). It has become one of the most widely used analytical tools among ecologists, with increasing prevalence in ecological research (Gao et al., 2025; Lai et al., 2019). Existing R packages address a range of biodiversity-related topics with applications to conservation, from microbial ecology (e.g., microeco, which is used to process environmental and microbial community data; Liu et al., 2021) to botany (e.g., bien, which centralizes multiple data sets in the Botanical Information and Ecology Network database; Maitner et al., 2018) to general biodiversity assessments (e.g., adiv, which complements existing R packages to provide a wide range of biodiversity indices; Pavoine, 2020). Despite the wide range of R packages available for ecological studies (Grainger & Gray, 2024), no single tool focuses on assessing the combined effects of roads and human footprints on species-level.

Here, we present ecoTolerance, a package tailored for systematic and streamlined calculation of RTI and HFTI and, by extension, derivative tolerance metrics, on datasets ranging from local community plots to continental-, and global-scale species inventories. The package offers automated data cleaning while its flexible input design allows users to combine global raster or local shapefiles, enabling analyses that range from a single protected area to the global scale. The functions return multi-level outputs (species, population and community-level summaries), facilitating direct comparisons of tolerance spectra within communities or among populations exposed to different anthropogenic influence.

By automating key steps, our package reduces coding time, allowing researchers to focus on hypothesis formulation, ecological inference, and decision-support.

Moreover, its versatility invites applications beyond global assessments: from comparing different populations across biomes to exploring how anthropogenic change reshapes functional redundancy across heterogeneous landscape mosaics by applying the species-specific tolerance indices.

Here we describe the workflow and the main functions of ecoTolerance and illustrate its practical utility through applied examples. The package is available on CRAN (https://cran.r-project.org/package=ecoTolerance). As an R package, it is open-source, fostering scientific transparency by making all codes available to be freely read and checked.

## 2 METHODS

### 2.1. Road Tolerance Index

To create a Road Tolerance Index (RTI), we used geographical coordinates based on species presence data to extract the distance between each record and the nearest road, and a road network vector. Considering the distance, at which the effect of a road can extend from its margin (Forman et al., 2003; Hamer et al., 2021; Torres et al., 2011), and the dispersal capacity of the focal taxa, ecoTolerance applies a reference distance that approximates a “zero-effect” threshold – the distance, beyond which road influence is assumed to be negligible. This threshold is set to 3.5 km by default, but users can adjust it to suit the ecology and biology of the focal taxa. Shorter distances indicate that the individual is potentially located in areas with some degree of impact by the road. Therefore, based on each individual’s distance to the nearest road, we calculate the Individual Road Index (RI_*i*_) following:

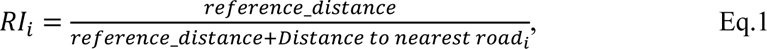

Where *i* represents each observation. This normalization ensures that the index ranges between 0 and 1. Each species has a distribution of densities of observations across the Individual Road Index. The median value of this distribution represents the Road Tolerance Index (RTI) for the species.

High RTI values (closer to 1) indicate that the species normally occurs near roads, reflecting tolerance, while low values (closer to 0) indicate that the species tends to occur in areas without roads, potentially reflecting low tolerance.

### 2.2. Human Footprint Tolerance Index (HFTI)

We constructed a continuous index to measure the tolerance of species to human-altered habitats using the global record of terrestrial Human Footprint raster (Mu et al., 2022). The global Human Footprint dataset integrates eight variables (built environments, population density, ALAN, crops, pasture, roads, railways, and navigable waterways) to create an index that reflects human pressures from different aspects and ranges from 0 to 50, where values < 1 indicate wilderness, < 4 indicates intact areas and 4 < indicate modified areas, potentially reaching 50 for highly modified areas (Mu et al., 2022).

Based on the geographical coordinates of each record (considered to be an individual), we extract the human footprint value within a 1-km buffer of each location, and rescaled it as following:

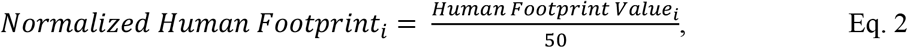

where *i* represents each observation and the scaling factor ensures the index ranges between 0 and 1. Each species had a unique distribution of densities of observations across the rescaled Human Footprint Index. The species-level Human Footprint Tolerance Index (HFTI) is based on the median value of this distribution.

A high HFTI (closer to 1) indicates that the species typically occurs in areas under high human impact, reflecting tolerance to human modification, while values closer to 0 suggest that the species is frequents areas with small or no human disturbance, reflecting low tolerance to human-altered habitats.

### 2.3. Overview of the ecoTolerance package

The ecoTolerance package was developed in the R programming language and calculates two indices - distance to roads and human footprint values, based on presence-only species occurrence data (R Core Team, 2025). The package provides a structured workflow (Figure 1) and consists of the core functions:

**FIGURE 1:**
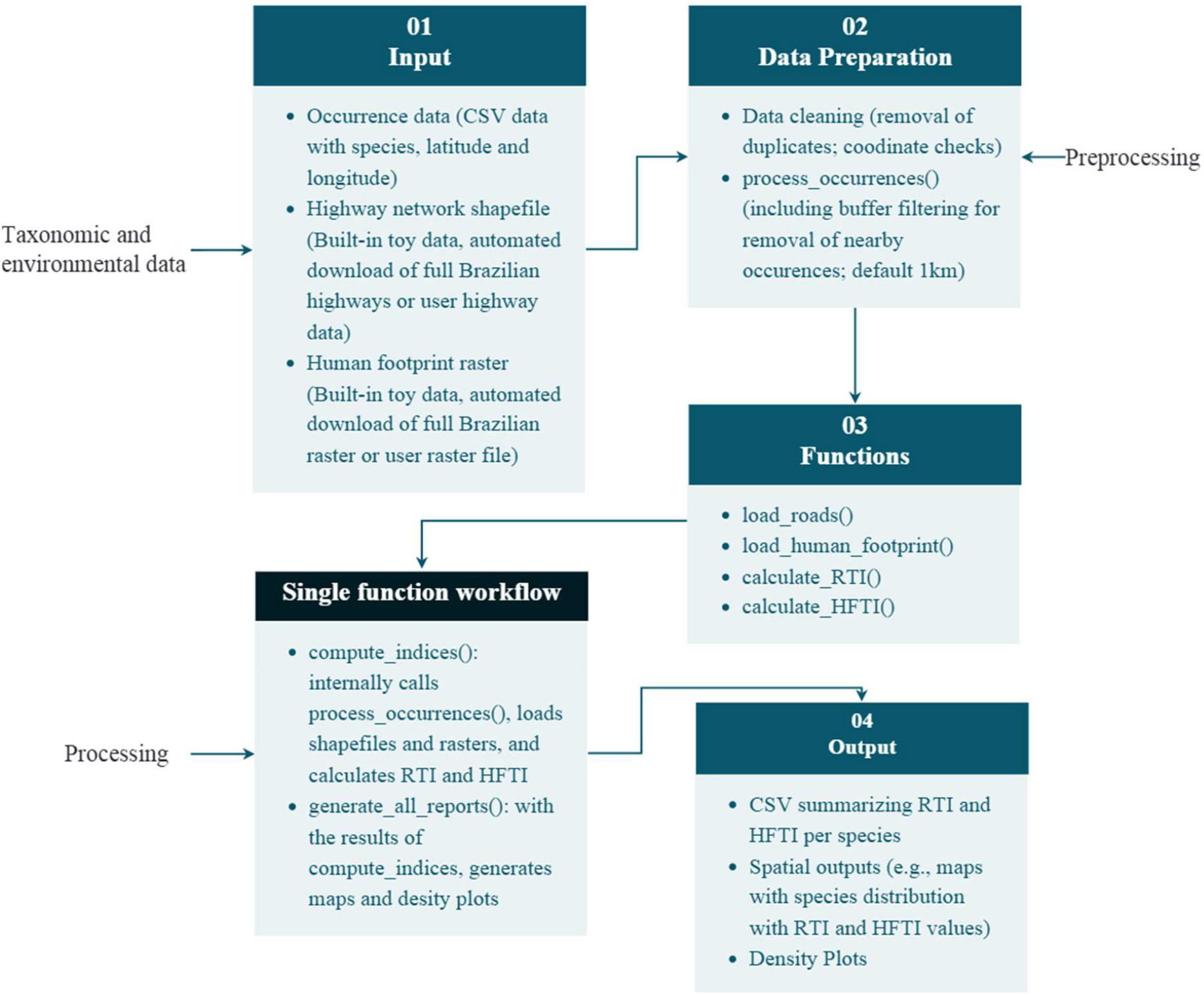
Flowchart of the processing performed in ecoTolerance

1. process_occurrences: cleans and filters species-occurrence records by (i) removing duplicates (multiple records of the same species with identical coordinates) and (ii) applying a buffer-based step that prevents points of the same species located within a user-defined radius (default = 1 km) from being over-represented.
2. load_roads and load_human_footprint: load the spatial layers required for the indices. By setting the argument *type = “full”*, the full Brazilian datasets (road-network shape file, CRS EPSG:4326), and the human-footprint raster layer provided by Mu et al. (2022)).
3. calculate_RTI: computes the Individual Road Index for each individual record based on Eq. 1, where dist is the minimal distance from the location of the species observation to the nearest road, and ref_dist is a user-defined reference distance based on a threshold distance to zero-effect (default = 3.5 km). This function also calculates the species Road Tolerance Index (RTI) based on the median RI_i_ for each species. RTI values range from 0 (very low tolerance) to 1 (high tolerance);
4. calculate_HFTI: calculates the Normalized Human Footprint Index (NHFI*i*) for each individual record based on Eq. 2, where the Individual Human Footprint Value is the average value of human footprint within a 1-km buffer zone around the location of the observation, rescaled to a 0-1 range. This function also calculates the Human Footprint Tolerance Index (HFTI) based on the median NHFI_*i*_ for each species;
5. compute_indices: a complete pipeline function that processes occurrence data, loads spatial layers, and calculates both indices in a single command. By adjusting the *data_type* argument, the function manages the automated download and integration of the full Brazilian spatial layers. It returns a list containing the processed data and the median RTI and HFTI per species.
6. generate_all_reports(): generates per-species maps and density plots of RTI and HFTI, and a CSV file with the two indices.

The package depends on *sf* (Pebesma, 2016) for handling vector data (roads and occurrence points), *raster* (Hijmans, 2010) for the human footprint raster, *dplyr* (Wickham et al., 2014) for data manipulation and *ggplot2* (Wickham, 2011a) for generating maps and graphs.

## 3 RESULTS

### 3.1. Implementation

We evaluated ecoTolerance’s performance and flexibility by testing it with a range of real-world datasets, spanning small (250 occurrences) to large scales (35,000 occurrences). The package successfully handled a test dataset of 35,000 occurrence points, processing both road distance calculations and footprint extraction within manageable times. Each core function (e.g., calculate_RTI, calculate_HFTI) can be run independently for users who only need partial functionality, and the single-function workflow (compute_indices) proved efficient, reducing manual scripting steps and lowering the potential for user error.

Because ecoTolerance relies on standard spatial classes, outputs can be readily input to other R packages, such as ggplot2 (Wickham, 2011b), for plotting. Our tests confirmed efficiency, reproducibility, and compatibility of ecoTolerance across datasets of varying size, making it suitable for both targeted studies (specialized datasets) and large-scale biodiversity analyses.

### 3.2. Spatial outputs

ecoTolerance returns a simple feature (*sf*) object (res$processed_data) containing the geometry of each occurrence along with RTI_value and HFTI_value columns. By using the generate_all_reports() function, and importing the shapefile of the study area, users can easily generate distribution maps scaled by the RTI and HFTI values, and density distribution plots for both indices.

These structures make it possible to correlate RTI and HFTI with species traits, habitat preference, or extinction risk categories. This tool can also elucidate the variability of species tolerance in different spatial-temporal contexts.

## 4 APPLICATIONS OF ecoTolerance

To demonstrate the versatility of *ecoTolerance*, we conducted three tests with distinct taxonomic groups (mammals, amphibians, and plants). Species-occurrence records were retrieved from GBIF via *rgbif* (Chamberlain et al., 2012) and cleaned using *CoordinateCleaner* (Zizka et al., 2019) to remove dubious points located at capitals, centroids, obvious outliers, or marine coordinates. Indices were computed with *compute_indices()*, and density plots and distribution maps were generated using *generate_all_reports()*.

For amphibians and mammals, we retained the default reference distance of 3.5 km – reflecting the typical road-effect zones and dispersal capacities, whereas for *Copaifera langsdorffii* we adopted a 1.5-km threshold, consistent with the estimates of road influence for arboreal plants (Forman et al., 2003; Hamer et al., 2021; Semlitsch, 2008; Smith & Green, 2005; Torres et al., 2011).

To demonstrate the use of ecoTolerance with mammals, we used *Silvilagus brasiliensis* (n = 274), a cursorial species, and *Bradypus variegatus* (n = 724), a slow-moving arboreal species. Despite contrasting life-history traits, both showed moderate road tolerance (RTI = 0.599 and 0.520, respectively) and comparable Human Footprint tolerance (HFTI = 0.279 and 0.341, respectively) (Figure 2). These results may suggest that even species with limited mobility can persist near roads when suitable arboreal cover remains, but both taxa still experience measurable constraints under higher human footprint values.

**FIGURE 2:**
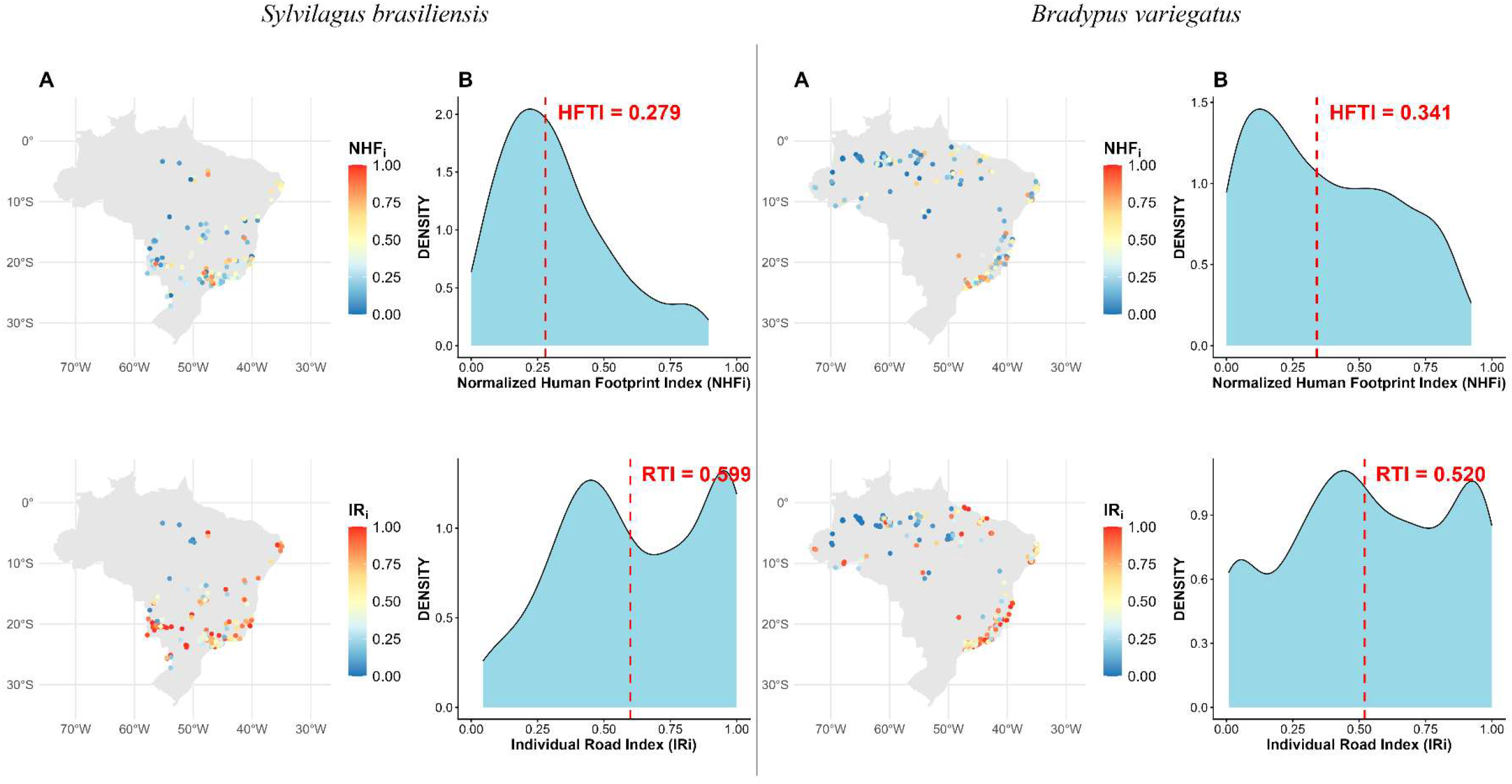
(A) Spatial distribution of occurrence records across Brazil colored by the normalized Human-Footprint Index (NHFi upper map) and by the Individual Road Index (IRi, lower map); warmer colors indicate higher tolerance values. (B) Kernel-density curves of NHFi (top) and IRi (bottom); red dashed lines mark the species-level medians (HFTI and RTI).

Using roadkill records (Miranda & Forti, in preparation) as an *a priori* indicator of vulnerability, we tested *Boana faber* (n = 1226) with 183 roadkill records and *Boana boans* (n = 151) with fewer than five collisions recorded. *B. faber* exhibited substantially higher tolerance (RTI = 0.654; HFTI = 0.392) than *B. boans* (RTI = 0.183; HFTI = 0.111) (Figure 3). The spatial overlap between *B. faber* occurrences and dense road networks can be used to support the utility of RTI as a rapid screening metric for road-impact susceptibility.

**FIGURE 3:**
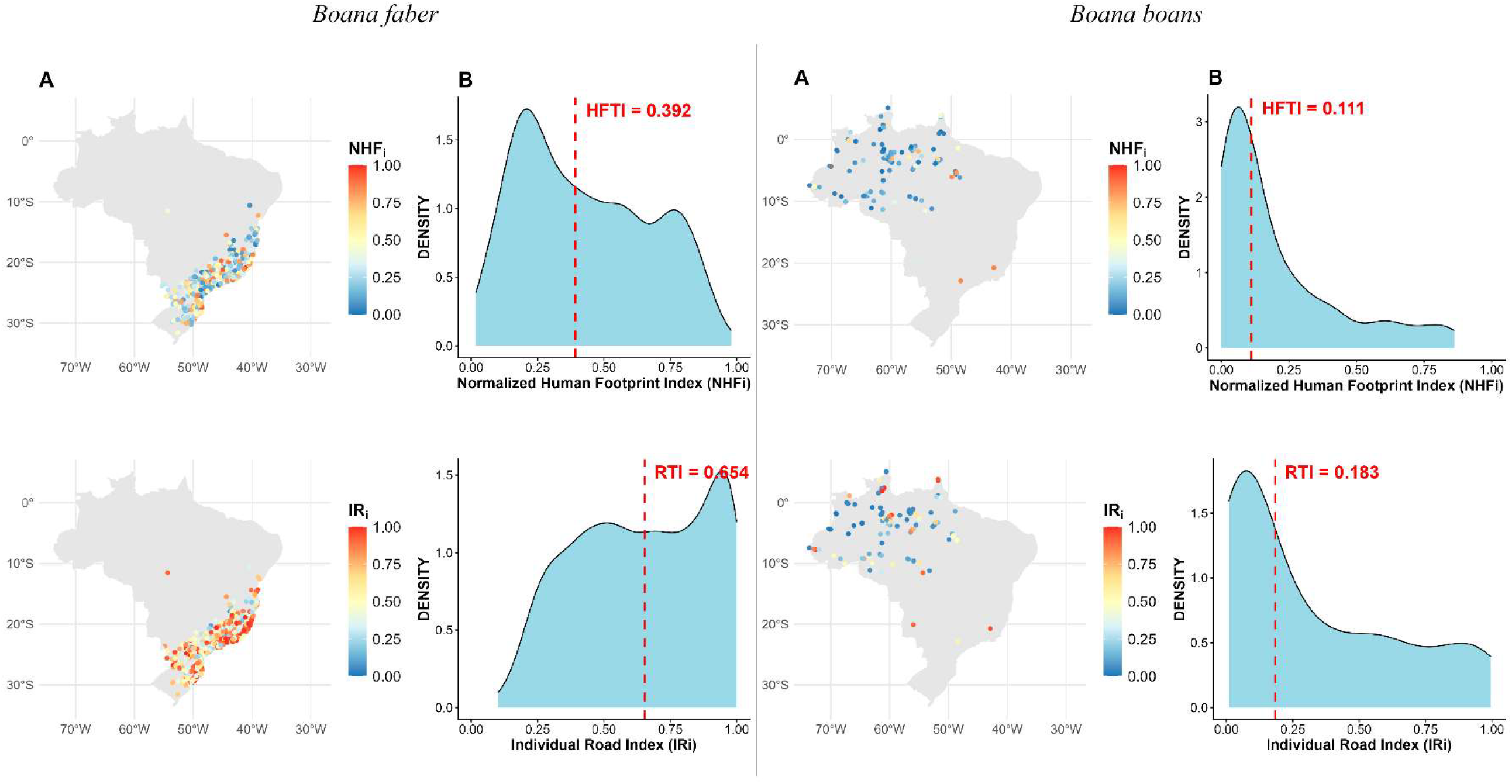
(A) Spatial distribution of occurrence records across Brazil colored by the normalized Human-Footprint Index (NHFi upper map) and by the Individual Road Index (IRi, lower map); warmer colors indicate higher tolerance values. (B) Kernel-density curves of NHFi (top) and IRi (bottom); red dashed lines mark the species-level medians (HFTI and RTI).

To test the applications with populations of a single species, we partitioned *Copaifera langsdorffii* into occurrence records in the Brazilian Cerrado (n = 924) and Brazilian Atlantic Forest (n = 483). Indices between biomes (RTI = 0.321 and 0.419 for the Cerrado and the Atlantic Forest, respectively, and HFTI = 0.248 and 0.348, respectively) indicate that conspecific populations can vary markedly in their tolerance profiles among biomes (Figure 4), likely reflecting contrasting land-use histories and road densities in these regions. The package thus facilitates population-specific conservation assessments and hypothesis testing with regard to local adaptation to anthropogenic pressures.

**FIGURE 4:**
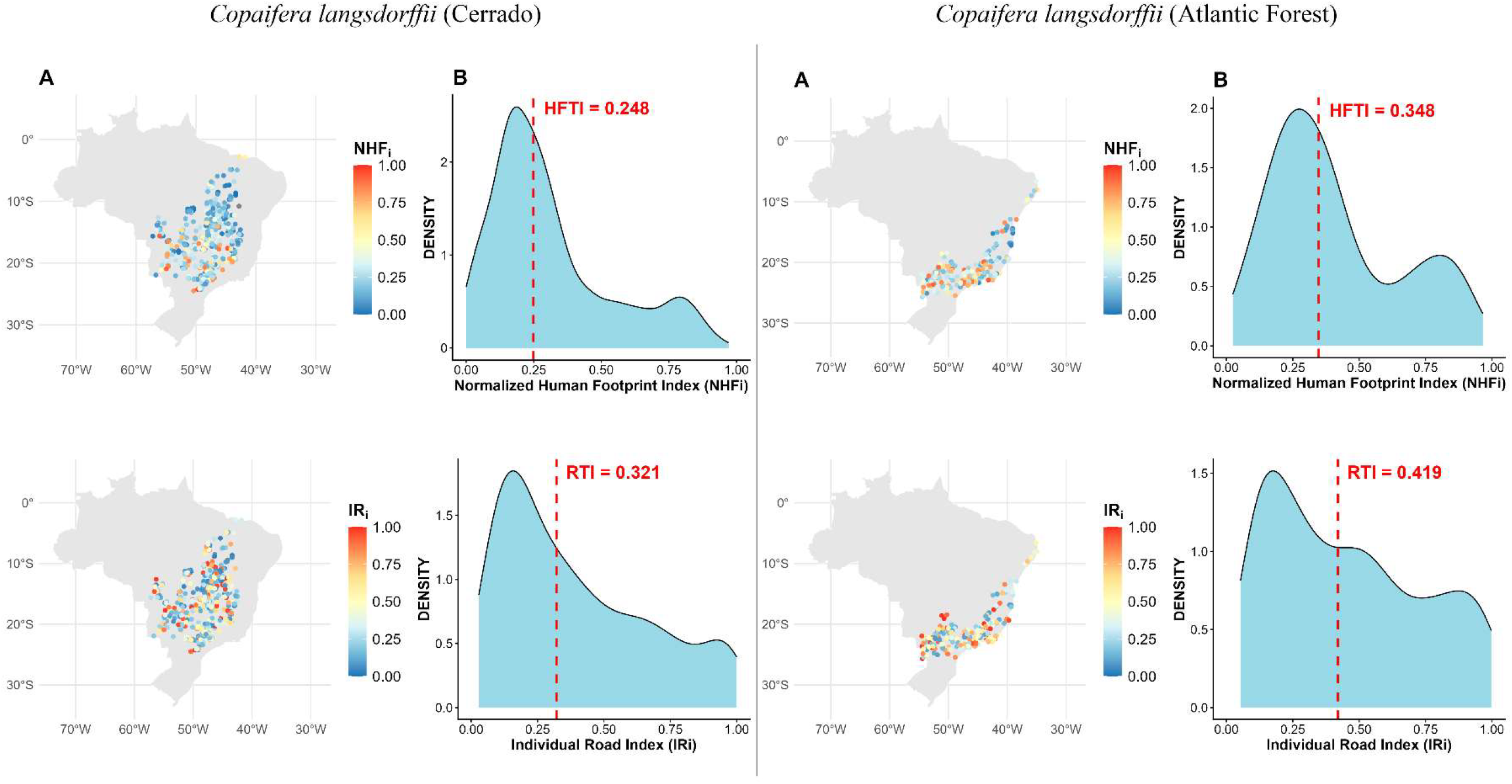
(A) Spatial distribution of occurrence records across Brazil colored by the normalized Human-Footprint Index (NHFi upper map) and by the Individual Road Index (IRi, lower map); warmer colors indicate higher tolerance values. (B) Kernel-density curves of NHFi (top) and IRi (bottom); red dashed lines mark the species-level medians (HFTI and RTI).

## 5 DISCUSSION

The ecoTolerance package provides the framework to quantify road tolerance and human footprint in species databases at different scales. By automating data cleaning and standardizing spatial calculations, ecoTolerance minimizes the risk of inconsistencies or manual errors. The package can be integrated into broader conservation studies, helping to identify priority areas for risk mitigation measures or planning for land-use changes, focusing on maintaining essential habitats and avoiding population declines of vulnerable species (Button & Borzée, 2021). By offering reproducible functions in R, ecoTolerance aligns with the need to establish open, transparent, and repeatable scientific practices that produce quality scientific knowledge in less time (Lowndes et al., 2017). In addition, the package allows uses to quickly adjust parameters (e.g., buffer sizes and reference distances) to match the ecological nuances of different taxa.

A possible restriction on using ecoTolerance in conservation studies is the requirement for precise spatial data, like georeferenced species location. If these data are not exact, the indices might not reflect the real tolerance of the species in question, which could lower the effectiveness of the model (Graham et al., 2008). In addition, future versions of ecoTolerance may incorporate automated checks for data quality related to roads and human footprint not initially included in the package and the combined use of additional layers related to metrics such as deforestation and fragmentation.

## ACKNOWLEDGEMENTS

We thank Judit K. Szabo for editing and reviewing the first draft. D. Miranda is grateful to CAPES for his PhD scholarship. L. R. Forti is supported by the Financiadora de Estudos e Projetos (FINEP) (grant number 01.23.0702.00).

## DATA AVAILABILITY

All data and supplemental material are available at https://doi.org/10.6084/m9.figshare.31424699

